# PU.1 Shapes Host Epigenetic Responses to Mycobacterium tuberculosis and Represents a Target for Host Directed Therapy

**DOI:** 10.1101/2025.09.09.675072

**Authors:** Jeremie Poschmann, Liu Jie, Caio Bomfim, Alice Mollé, Wenjie Sun, Taylor W. Foreman, Colette Cruesy, Gosset Pierre, Deepak Kaushal, Gennaro de Libero, Celine Cougoule, Shyam Prabhakar, Amit Singhal

## Abstract

*Mycobacterium tuberculosis* (*Mtb*) infection induces widespread epigenetic changes in monocytes and macrophages. To characterize the critical factor responsible for *Mtb*-induced epigenetic responses, we performed an integrative analyses of chromatin modification (H3K27ac), transcription factor binding, and gene expression in human monocytes and macrophages. Our data identified PU.1 (SPI1) as a key transcriptional regulator to be upregulated and bound to gene promoters in response to *Mtb*. PU.1 enrichment was associated with enhanced proinflammatory and anti-apoptotic gene programs. PU.1 expression was also elevated in the lung tissues from *Mtb*-infected macaques and tuberculosis patients. PU.1 knockdown enhanced human macrophage apoptosis, dampened inflammation and reduced bacterial survival, suggesting its functional role in supporting *Mtb* infection. Pharmacological inhibition of PU.1 replicated the effects of knockdown, restricting *Mtb* growth without cytotoxicity. These findings identify PU.1 as a central node in the host epigenetic response to *Mtb* and a candidate for host-directed therapeutic targeting in tuberculosis.

## Introduction

Pathogens have evolved strategies to promote their survival by regulating the host immune response via modulating the epigenetic landscape of immune cells (Fol et al., 2020; Silmon de Monerri and Kim, 2014). These infection-induced epigenetic changes, which are plastic and dynamic in nature, have a direct impact on the gene expression in host cells and the pathogenesis of the diseases (Fischer, 2020; Russ et al., 2014).

*Mycobacterium tuberculosis* (*Mtb*), the causative agent of tuberculosis (TB), have also demonstrated the ability to reprogram and disable key antibacterial response pathways, many of which are governed by epigenetic mechanisms regulating the transcriptional machinery (Del Rosario et al., 2022; Khadela et al., 2022; Madden et al., 2022; Singh and Nagaraja, 2025). We and others have demonstrated perturbed epigenetic mechanisms, involving both histone modifications and DNA hypermethylation, in the blood cells of TB patients (Del Rosario et al., 2022; DiNardo et al., 2020). However, the host factor(s) mediating the *Mtb*-induced epigenetic and down-stream transcriptional changes are still not known. Such epigenetic driver could serve as targets for host-directed therapy (HDT) strategies in TB (Dai et al., 2024; Gai et al., 2023).

Here, using histone H3 acetylation at lysine 27 (H3K27ac) and chromatin immunoprecipitation with massively parallel DNA sequencing (ChIP-seq) analysis we report that *SPI1* / PU.1, an ETS family transcription factor essential for myeloid cell differentiation (Kastner and Chan, 2008), serves as a master regulator of macrophage transcriptional and apoptotic program in response to *Mtb* infection. Using small molecules that selectively inhibit PU.1 DNA binding we show that PU.1 could be therapeutically targeted to improve *Mtb* control via enhancing apoptosis response as part of a host-directed strategy against TB. Our study thus identifies PU.1 as a modifiable epigenome regulator to improve bacterial control through host cell reprogramming of macrophages.

## Results

### *Mtb* infection causes genome-wide epigenetic changes in host cells

In order to investigate potential epigenetic alterations that occur during *Mtb* infection, we set out to estimate genome-wide H3K27ac, a histone mark that is known to be enriched at active promoters and enhancers (del Rosario et al., 2015) and to be highly correlated with gene expression (Chaumette et al., 2022). *Mtb* infected and uninfected THP-1 cells were collected in parallel and H3K27ac chromatin immunoprecipitation followed by sequencing (ChIP-seq) profiles were generated at the following time points of infection (0h, 24h, 48h and 72h) (see Methods). On average, we sequenced 10-23 million non-redundant mapped reads per sample and time point. To estimate the success and the utility of H3K27ac ChIP-seq to efficiently monitor changes during *Mtb* infection we investigated whether infection leads to changes in H3K27ac enrichment at key genes of immune responses (*IFIT* genes, *CTSG* and *CXCR2*) (Figure 1A). Indeed, acetylation of the *IFIT 1, 2* and *3* loci increased in the *Mtb* infected samples, suggesting a persistent epigenetic activation of these genes (median fold-change 4.4, 2.1 and 5.7, respectively). The converse was true for the *CTSG* and *CXCR2* genes, which displayed a reduction of their acetylation in infected samples compared to control (median fold-change 0.3 and 0.2).

**Figure 1.**
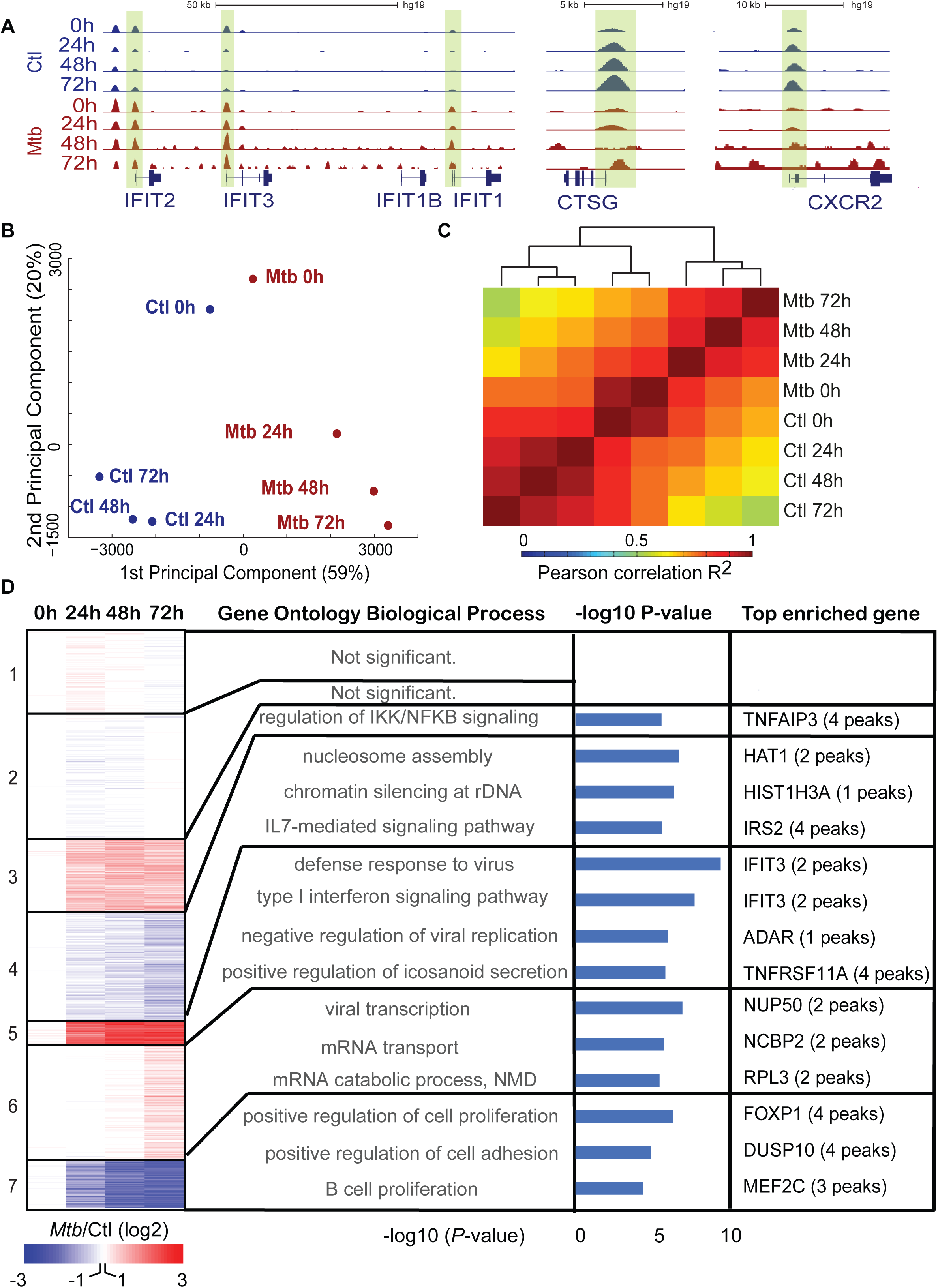
*Mtb* infection reprograms H3K27ac in THP-1 cells on a genome-wide scale. (A) Genome browser tracks of H3K27ac among control (Ctl, in blue) and *Mtb*-infected THP-1 (*Mtb*, in red) cells at 0h, 24h, 48h and 72h of infection at the representative chromosome loci *IFIT2, IFIT3, IFIT1, CTSG, CXCR2*. Promoter peaks are shaded for each gene. (B) Principal component analysis of H3K27ac across time and condition. (C) Heatmap showing clustering of pair-wise Pearson correlation R^2^ of H3K27ac data. (D) K-means clustering of H3K27ac fold change between control and *Mtb* samples at each time point. Gene ontology (GO, biological process) has been indicated for each cluster with the −log_10_ *P*-value. In addition, the top enriched genes for H3K27ac peaks are shown for each GO.

To quantify epigenetic changes more globally, we examined differential acetylation between infected and control samples based on the above-described chromatin state maps. Principal components analysis (PCA) of H3K27ac peaks revealed that a genome-wide epigenetic response to infection varied over time (Figure 1B). Notably, the first principal component (59%) clearly separated the infected *vs* control (Ctrl) samples. The second principal component (20%) reflected temporal dynamics after contact with *Mtb*. After the first 24 h of infection with *Mtb*, the non-infected samples showed less temporal variability than the infected samples. We hypothesized that the former samples responded to the change in conditions during the first 24 hours and then settled into a stable state, whereas the latter continued to respond to the bacterial infection. These inferences are further supported by hierarchical clustering of the samples based on pair-wise Pearson correlations of H3K27ac peak heights (Figure 1C). Notably, three clusters are visible: the 0h time points of infected and control samples, a control cluster and an *Mtb* infected cluster. Collectively these data indicate that *Mtb* infection triggers a global and dynamic epigenetic response in the host cells, which extends up to 72h.

We next identified the molecular pathways underlying the epigenetic changes. For this, we first calculated the infection response time course of each regulatory element (H3K27ac fold change vs time). We then used k-means clustering of these epigenetic trajectories to detect 7 peak clusters (Figure 1D, Supplemental Table 1). Two of these clusters (clusters 3 and 5) were upregulated from 24-72h and another 2 were down regulated (clusters 4 and 7). Gene ontology (GO) analysis of the upregulated clusters showed GO enrichment for the nuclear factor kappa B (NF-κB) signaling, type 1 IFN signaling, defense response to virus and other immune responses (Supplemental Table 2). In contrast, the downregulated clusters were enriched for nucleosome assembly, chromatin silencing, regulation of cell proliferation and cell adhesion, and B cell proliferation. In addition, Cluster 6 showed moderate up-regulation at the final time point (72h) associated to viral transcription and mRNA metabolism. Figure 1D depicts the genes associated with the largest number of acetylation peaks for each cluster for each significant GO term. Of note, transcriptomic and epigenetic data have previously shown to activate similar pathways upon *Mtb* infection in monocytes and macrophages (Del Rosario et al., 2022; Madden et al., 2023; Olson et al., 2021; Poladian et al., 2023), thus qualitatively validating the use of histone acetylation as a readout for molecular immune responses.

### Discovery of transcription factors that mediate the H3K27ac response

Genomic regions marked by H3K27ac peaks are bound by transcription factors (TF) that recruit co-factors and mediate transcriptional regulation (Sun et al., 2016). Thus, H3K27ac rich chromatin locations should contain an excess of high-affinity binding sites for master TFs that mediate the immune response to *Mtb* infection. We therefore performed TF motif enrichment analysis on each of the up- and down-regulated H3K27ac clusters (Clusters 3-7; Figure 2A). The most enriched motifs (see Methods, Supplementary Table 3) matched well-known mediators of immune response to *Mtb* infection, such as IRF8, IRF7, NFKB, STAT1, ATF and JUN (Esquivel-Solis et al., 2009; Fallahi-Sichani et al., 2012; Kumar et al., 2020; Madden et al., 2023; Marquis et al., 2011). In addition, we found strong and consistent enrichment of PU.1 motifs in three out of five clusters.

**Figure 2.**
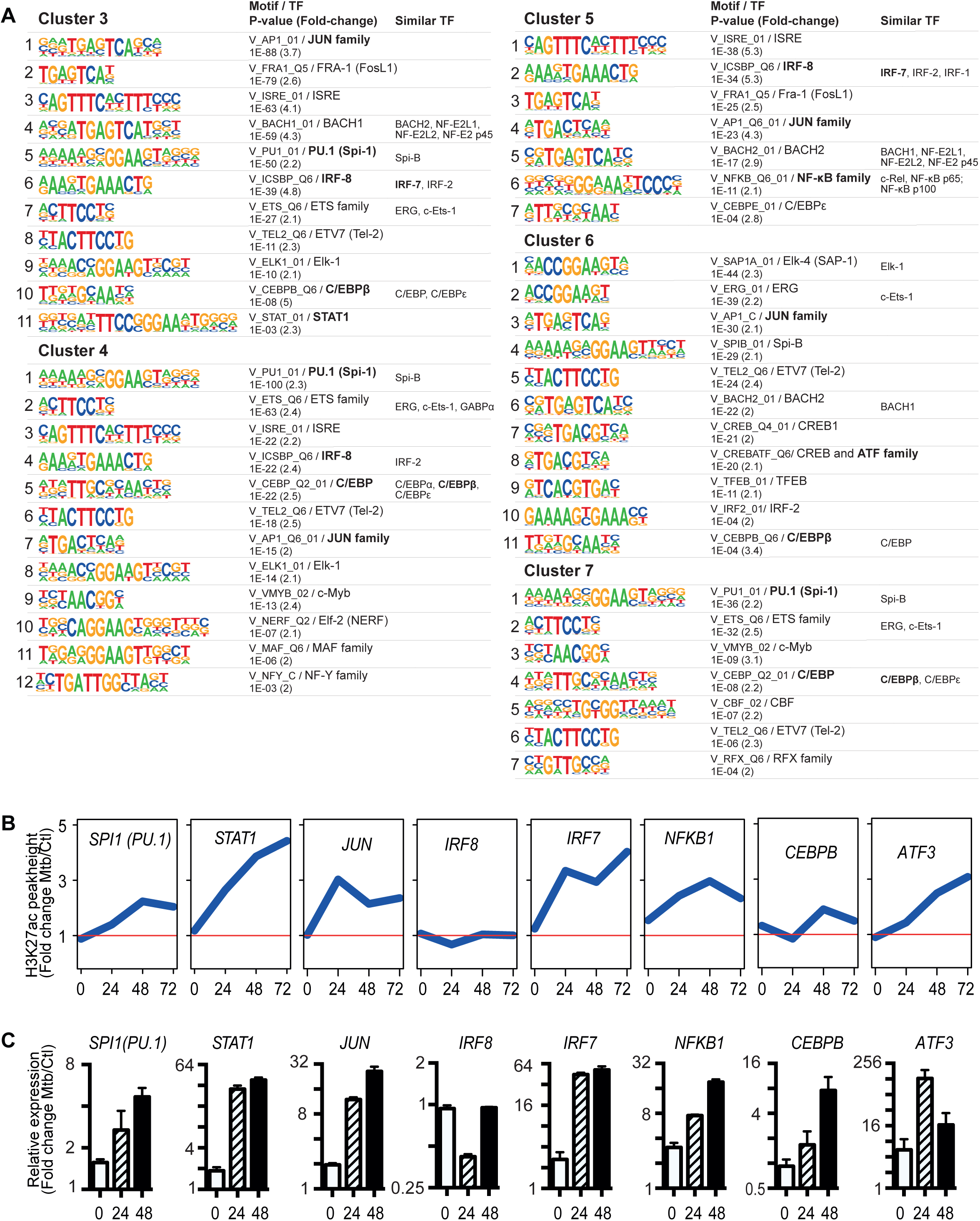
Identification of transcription factors (TF) implicated in the response to *Mtb* infection. (A) A representative transcription factor of the subfamily identified in each cluster (cluster 3-7, Figure 1D) has been depicted. Shown is the motif, the *P*-value, the enrichment fold change, the corresponding TF subfamilies and other TFs of the same subfamily identified. (B). The normalized H3K27ac peak heights of TF identified in panel A around their promoter regions. The peak height differences between *Mtb* and control (Ctl) cells is shown across time (0, 24, 48 and 72h) for each promoter. (C) Relative gene expression (expression of gene in *Mtb* infected cells relative to ctrl cells) of TFs quantified by RT-qPCR during *Mtb* infection at 0h 24h and 48h.

PU.1 has been described as a master regulator of immune cell fate determination (Singh, 2008; Spooner et al., 2009), and has been associated with (i) the macrophage enhancer activity in response to LPS stimulation (Ghisletti et al., 2010), and (ii) the induction of allele-specific IL-1β expression in TB (Zhang et al., 2014). To further corroborate the motif enrichment, we examined histone acetylation at the promoters of the same TFs over time (Figure 2B). Indeed, promoter acetylation increased over time (positive slope) for 7 of the 8 TFs, except IRF8. This motif and promoter acetylation analysis of TFs was supported by *Mtb*-infection induced gene expression trajectories of all TFs, including PU.1/*SPI1* (Figure 2C). Together, the above motif enrichment and gene regulation results suggest that PU.1 may play an unexpectedly broad role in orchestrating the epigenetic dynamics and transcriptional programs of *Mtb*-infected cells, extending beyond its established function in lineage specification. We therefore prioritized PU.1 for further investigation.

### PU.1 binding regulates the immune response to *Mtb* infection

Since the mRNA level of *SPI1* (PU.1) was robustly upregulated upon *Mtb* infection, we also examined PU.1 protein expression (Figure 3A). Consistently with the mRNA levels, PU.1 was upregulated at least until 48 h post-infection (Figure 3A). We next interrogated the role of PU.1 in the *Mtb* immune response, by performing ChIP-seq of PU.1 in control and *Mtb*-infected cells. A total of 66,968 binding sites across the 6 conditions (control and infected, three time points), of which 23,009 overlapped H3K27ac peaks was observed (Methods, Figure 3B). PU.1 binding was substantially stronger when it coincided with histone acetylation (Supplemental figure 1A). Moreover, relative to non-acetylated PU.1 binding sites, PU.1 sites in acetylated regions were three-fold more likely to bind promoters (Figure 3C). We therefore focused our attention on the 23,009 PU.1 binding sites that lie within H3K27ac peaks, to investigate the mechanistic role of PU.1 in controlling the epigenetic dynamics of *Mtb* infection. We tested systematic association between PU.1 binding and histone acetylation within the 5’ regulatory element clusters that were dynamic in response to infection (Clusters 3-7, Figure 1). In all clusters, we found significant changes in PU.1 binding from 0 h to 24 h (Figure 3D). PU.1 binding was significantly elevated in all upregulated clusters (Clusters 3, 5 & 6) and attenuated in all downregulated clusters (Clusters 4 & 7), which is consistent with a role of transcription factors in recruiting histone acetyl-transferases to regulatory elements. Together, these results suggest that PU.1 functions as a master regulator of the global epigenetic response to intracellular *Mtb* infection.

**Figure 3.**
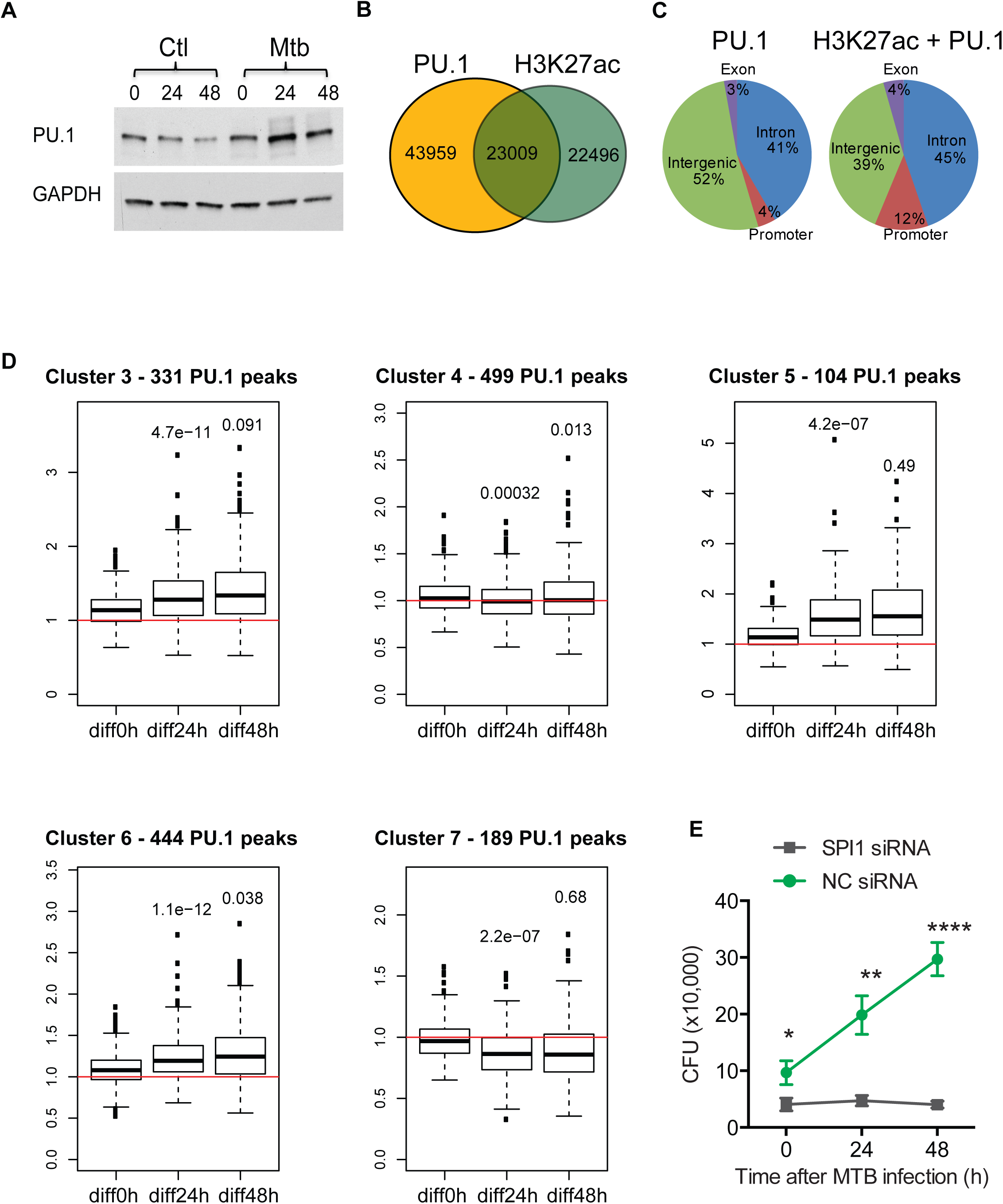
The transcription factor PU.1 plays a major role in the response to *Mtb* infection. (A) Protein expression of PU.1 in *Mtb* infected THP-1 cells at respective time points. GAPDH protein expression was used as a loading control. (B) Overlap of H3K27ac and PU.1 ChIP-seq peaks in THP-1 cells. (C) Comparison of genomic annotation of PU.1 peaks those lie within H3K27ac peaks (labelled as H3K27ac+PU.1) vs those that are not in H3K27ac peaks (labelled as PU.1). (D) Boxplots of PU.1 binding within H3K27ac peaks in the clusters identified in Figure 1D. Shown is the fold change of peak heights (y-axis) between *Mtb* infected and control cells at the indicated time points (0h, 24h and 48h). *P*-values of Wilcoxon rank sum test comparing the 0h to 24h or 48h are indicated in the boxplots. (E) CFU assay quantifying the *Mtb* growth in *SPI1* knockdown THP-1 cells vs scrambled cell (NC). Data in E is shown as Mean±SD; *P*-value, two-tailed Student *t*-test at each time point, *<0.05, ** <0.01, ***<0.001.

We next sought to understand the functional role of PU.1 expression and mycobacterial survival. Colony-forming unit (CFU) assay was performed on wild type and PU.1 knockdown THP-1 and human monocyte differentiated macrophages (hMDMs). Knockdown efficiency is shown in Supplemental figure 1B. PU.1 knockdown significantly suppressed the growth of *Mtb* and BCG (Figure 3E and Supplemental figure 1C). PU.1 knockdown did not affect proliferation of THP-1 cells (Supplemental figure 1D). Together, our data suggests that the upregulation of PU.1 in infected cells is beneficial to mycobacterial survival.

### In vivo characterization of PU.1 expression in active TB patients and *Mtb*-infected macaques

We next evaluated in vivo and clinical significance of *Mtb*-mediated PU.1 upregulation observed *in vitro* (Figure 3A). First, we assessed the *SPI1* mRNA expression profiles among peripheral blood of active pulmonary TB patients, latent TB individuals, and healthy controls from four different cohorts (Figure 4A-D, Supplemental table S4) (Berry et al., 2010; Cai et al., 2014; Kaforou et al., 2013). In all these cohorts’ *SPI1* mRNA levels were highest in active pulmonary TB patients compared with healthy and latent TB individuals (Figure 4A-C). In active pulmonary TB patients undergoing anti-TB chemotherapy *SPI1* mRNA in blood decreased after two months of therapy and reached levels comparable with those observed in latent TB individuals by the six months of treatment (Figure 4D).

**Figure 4.**
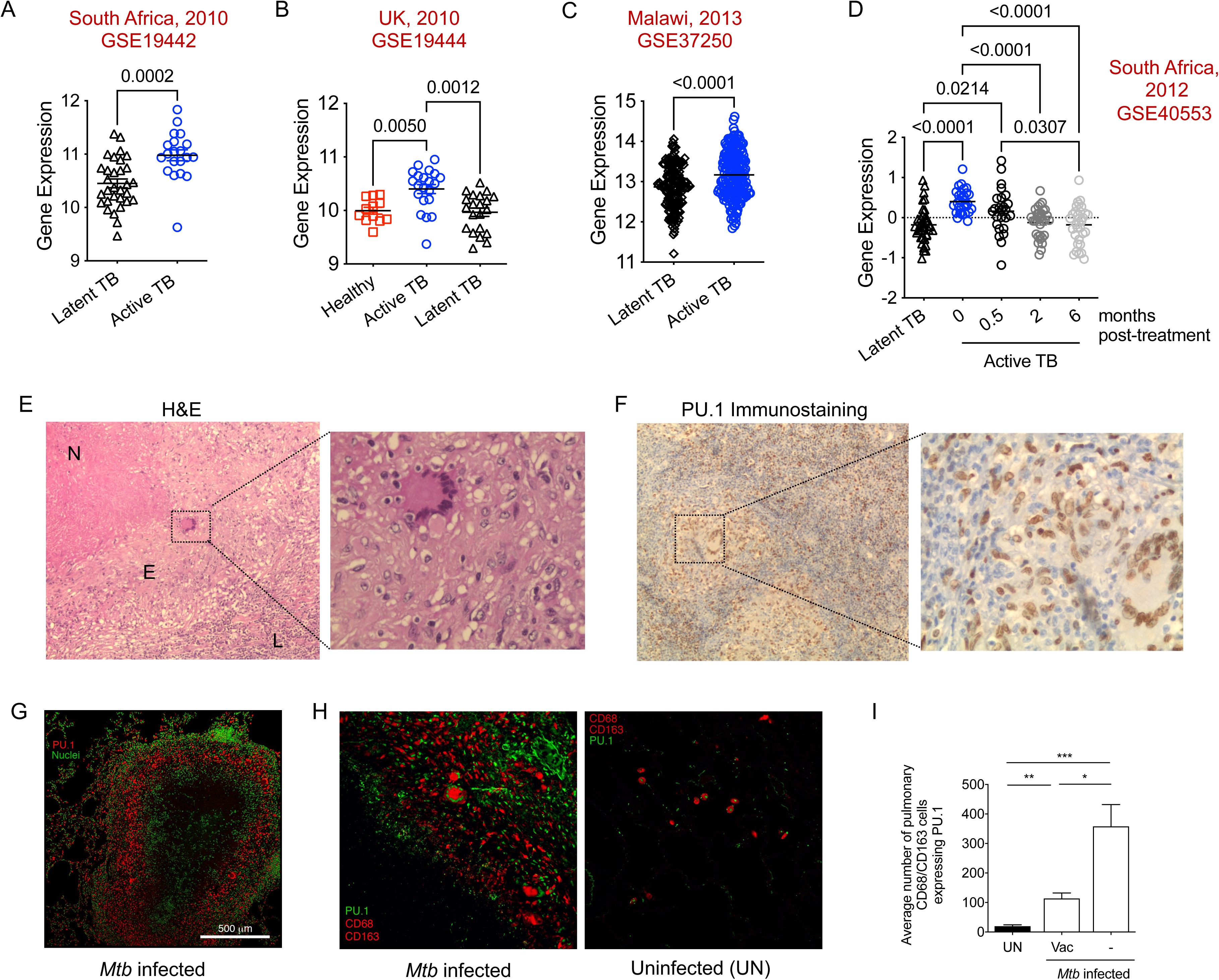
*Mtb* infection upregulates SPI1/PU.1 expression in TB patients and *Mtb*-infected macaques. (A-C) Raw intensity values for *SPI1* mRNA expression in the blood data set of different TB cohorts (Supplemental Table 4). Black bar indicates the median. \**P*-value by Mann-Whitney test. (D) *SPI1* mRNA expression data from active TB patients undergoing standard chemotherapy treatment from South Africa, 2012 cohort. Black bar indicates the median, \**P*-value by one-way ANOVA test. (E) H&E staining of granuloma in the cervical lymph node of an extrapulmonary TB patient. N - necrotic area, E - Epithelioid macrophages, L – Lymphocytes. Zoomed region of multinucleated giant cell is shown. (F) Immunostaining of granuloma in E with PU.1 antibody, showing strong nuclei staining of epithelioid macrophages. Weak staining was also observed in cytoplasm. (G) Immunostaining of lung tissue for PU.1 of a representative *Mtb*-infected macaque with active disease. Presence of increased PU.1 (red) in non-necrotic (N) area is shown. Nuclear staining is in green. Magnification 2×. (H) PU.1 (green) and CD163/CD68 (red) co-immunostaining in the lung tissue of uninfected and *Mtb*-infected macaque. Magnification 20×. (I) Similar to H immunostaining was performed on the lung tissue from uninfected; BCG-vaccinated- and unvaccinated-*Mtb*-infected macaques. Compiled data of percentage of pulmonary CD163^+^CD68^+^ cells expressing PU.1 is indicated (*n* = 8 – 20). *P*-value by one-way ANOVA test. * *P* < 0.05, \*\**P* < 0.01, \*\*\**P* < 0.001.

To investigate the expression of PU.1 in the tissues of *Mtb*-infected host, we first performed immunostaining in the lymph node sections from an extrapulmonary TB patients. H&E staining of the granuloma of lymph node TB subject shows well demarcated necrotic area, followed by epithelioid macrophages and an outer layer of lymphocytes (Figure 4E). Multinucleated giant cells, characteristic of human TB, were also observed. The nucleus of epithelioid macrophages and giant cells demonstrated a positive staining of PU.1 (Figure 4F). Of note, lymphocytes did not show PU.1 staining.

We further performed immunostaining on the lung sections from *Mtb-*infected macaques, which showed expression of PU.1 among leukocytes, mainly CD163^+^CD68^+^ macrophages (Figure 4G and H). Granulomas of BCG-vaccinated *Mtb-*infected macaques (Kaushal *et al*., 2015), which we earlier screened for SIRT expression (Cheng et al., 2017), showed lower levels of PU.1 among CD163^+^CD68^+^ macrophages as compared with unvaccinated *Mtb-*infected macaques (Figure 4I). Taken together, these data indicate an association of lower *SPI1*/PU.1 expression with protection from *Mtb* infection *in vivo*.

### Genome-wide identification of PU.1 targets in *Mtb* infection

We next aimed to characterize the functional consequence of PU.1 by identifying its downstream targets using a PU.1 knockdown strategy. In THP-1 cells, we profiled gene expressions comparing siRNA-mediated knockdown of PU.1 versus scrambled control siRNA (NC). This revealed 1,929 up- and 2,176 down-regulated genes whose expression was affected by PU.1 knockdown. In parallel, we performed the gene expression analysis in *Mtb* infected cells 24h post-infection, which identified 1,225 up- and 1,255 down-regulated genes compared to uninfected control (Ctrl) cells. The two datasets were merged, which yielded a total set of 2,480 differential expressed genes (DEGs). We compared the fold change of these DEGs, enabling us to identify two sets of genes that were regulated by PU.1, and were also modulated in response to *Mtb* (black circles, Figure 5A, Supplemental table 5). The first comprised 871 genes that were downregulated by PU.1 (viz. upregulated when PU.1 was knocked down) and concurrently downregulated during infection (upper left quadrant, Figure 5A). The second included 942 genes, which were upregulated by PU.1 (i.e., downregulated upon PU.1 knockdown) and upregulated in response to infection (lower right quadrant, Figure 5A). Thus, in total 1,813 (871+942) genes (73.1%) of the total 2,480 *Mtb*-responding genes were characterized as PU.1-regulated in our dataset.

**Figure 5.**
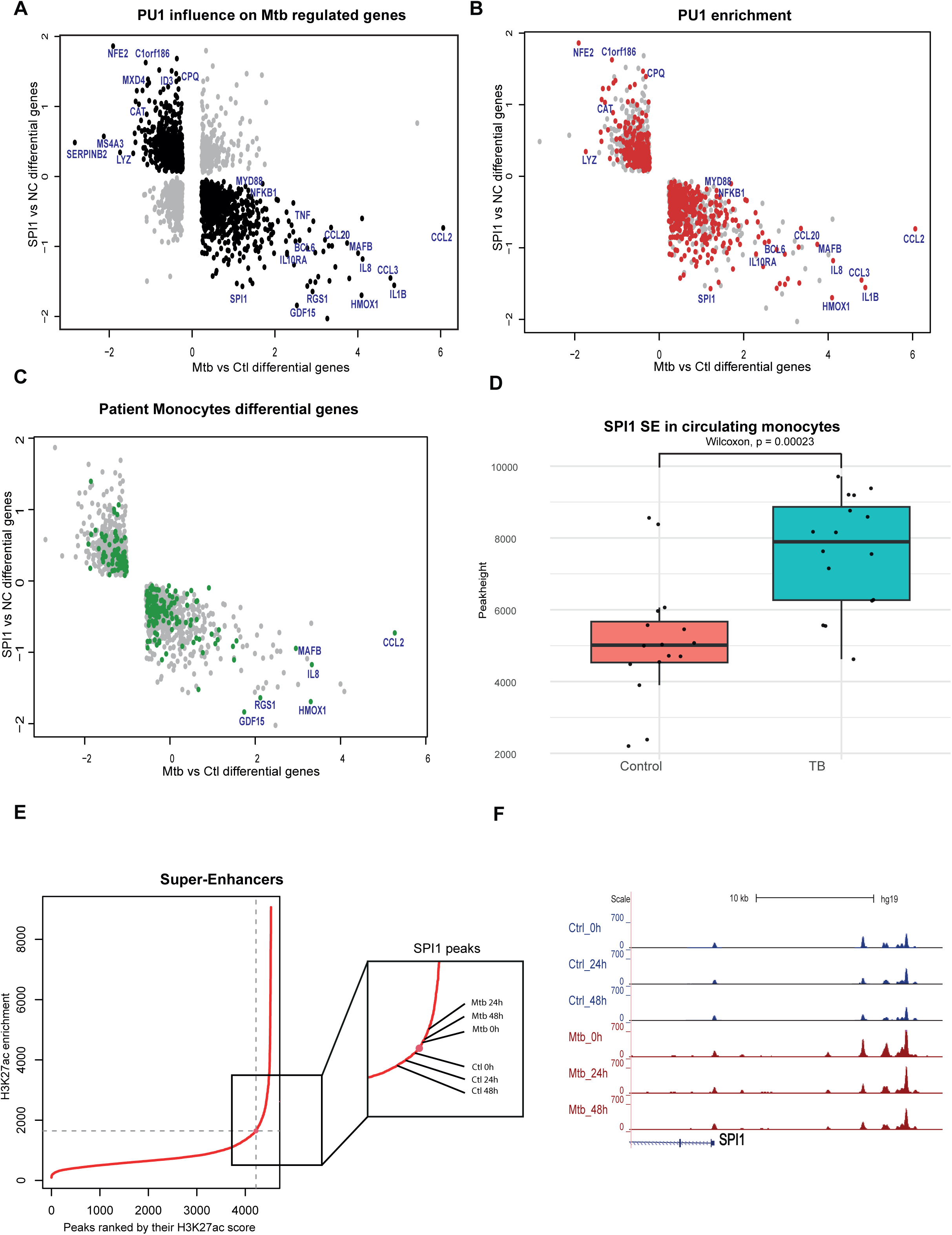
PU.1 is a major driver of the transcriptomics host response towards *Mtb* infection. (A) Scatterplot showing PU.1 regulated genes indicated as black dots. Genes not affected by PU.1 are shown in grey. X axis shows the log_2_ fold change of significantly differential expressed genes (DEGs) in response to *Mtb* infection at 24h in THP-1 cells. Y-axis shows the log_2_ fold change of DEGs in SPI1 knock down versus scrambled knockdown (NC) THP-1 cells. (B) Scatter plot in A showing PU.1 regulated genes in grey and genes which had PU.1 peaks at their promoter are labelled in red. (C) Same as panel B but genes were colored green if they were also differentially expressed in circulating monocytes from active TB patients. Gene expression data of TB monocytes was obtained from (Del Rosario et al., 2022). (D) Genome browser track of the PU.1 peaks at *SPI1* locus in *Mtb* infected and control THP-1 cells at indicated time points. (E) Super enhancers ranked by peakheight identified in the H3K27ac CHIP-seq dataset. The point of inflection (straight red line) indicated whether a peak is a super enhancer. The zoom-in shows the *SPI1* super enhancer peakheight for each condition (Ctl and *Mtb* at different time points). (F) Boxplot displaying the peakheights of the *SPI1* super enhancer in circulating monocytes of active TB patients and healthy control. H3K27ac data was obtained from (Del Rosario et al., 2022). *P*-value, Wilcoxon rank sum test.

Next, we examined whether PU.1 ChIP-seq peaks were near the 1,813 PU.1-regulated genes and found that 803 genes (44.3%) promoters were bound by PU.1 (red color genes, Figure 5B). Relative to the entire set of all 21,460 genes, the 1,813 PU.1-regulated genes were significantly enriched for PU.1 binding near the transcription start site (Fisher exact test, *P*-value = 2.2×10^−16^), indicating that PU.1 is a major driver of the host response to *Mtb* infection.

We next evaluated whether the *in vitro* regulated PU.1 genes were also differentially expressed in TB patients. For this, we used our recently reported transcriptomic data from the circulatory monocytes of healthy controls and active TB patients (Del Rosario et al., 2022). We mapped the 1,980 DEGs between healthy controls and active TB patients on the PU.1 regulated genes (Figure 5C). We found that 221 DEGs (out of 1,980) were also PU.1 regulated (Fisher exact test, *P*-value = 1.8×10^−4^) indicating a significant overlap between gene expression changes in circulating monocytes and upon *in vitro Mtb* infection. In summary, we identified a set of genes that are directly and indirectly regulated by PU.1 in response to *Mtb* infection.

### Extensive regulation of PU.1 super enhancer during *Mtb* infection

In our data, we noted that *SPI1* / PU.1 is differentially expressed in response to *Mtb* infection and was bound by itself (Figure 5B, lower right quadrant). We therefore verified in detail whether PU.1 is binding to its own gene *SPI1*. Indeed, multiple PU.1 peaks were prominent and increased in response to *Mtb* near *SPI1* gene (Figure 5D), indicating that PU.1 regulates itself in response to *Mtb* infection. Since multiple PU.1 binding sites were found surrounding the *SPI1* gene we carried out a super enhancer (SE) analysis to determine whether *SPI1* is regulated by a SE, which are large genomic regions containing multiple enhancers and are thought to control genes important in development, differentiation, and inflammatory response (Blayney et al., 2023; Higashijima and Kanki, 2022; Pott and Lieb, 2015). SEs were detected by aggregating H3K27ac peaks to generate SEs and measuring the sum of peakheights within the SEs (see Method). A curve was then drawn, ranking the SE by its peakheight (Figure 5E). The point of inflection of this curve determined whether a H3K27ac peak was a SE or not. Interestingly, we found that *SPI1* was right at the point of inflection at the 0h time point then its peakheight was increased in response to *Mtb* infection (24h, 48h & 72h), however remained decreased in the control conditions (see Figure 5E zoom-in). Thus, the combination of the increased binding of PU.1 and the increased acetylation of this PU.1-SE indicates that *SPI1* was a SE regulated by itself in response to *Mtb* infection. Since these results were derived from *in vitro* THP-1 data we analysed the SE state of *SPI1* using H3K27ac ChIP-seq in circulating monocytes from healthy controls and active TB patients, which we published earlier (Del Rosario et al., 2022). In this cohort dataset also our analysis identified *SPI1* as a SE in circulating monocytes, and found it to be significantly upregulated in the TB patients (Wilcoxon rank sum test, *P*-value = 2.3×10^-4^; Figure 5F, Supplemental table 5). Taken together, this indicated that PU.1 is a critical transcription factor extensively regulated at an epigenetic level in TB.

### Characterization of PU.1 target genes in response to *Mtb* infection

To ascertain the biological functions controlled by PU.1, we characterized functional enrichment within the 1,813 PU.1-regulated genes (red color genes, Figure 5B). Relative to the entire set of expressed genes, the PU.1-regulated genes were highly enriched for gene ontologies (GOs) such as defense response, apoptosis, proliferation, wounding, and immune responses (Figure 6A). Several of the PU.1-regulated genes was associated to one or more GOs (Figure 6B). For example, the most upregulated genes, *CCL2* and *IL1Β*, were associated with immune response, proliferation, and apoptosis. We verified if these two genes were also changed at the protein levels using ELISA assay in the supernatant after 24h post-infection comparing control vs *SPI1* siRNA knockdown cells (Figure 6C). We also checked whether PU.1 was directly binding in the vicinity of these two genes. For both genes the infection conditions revealed exclusive PU.1 binding close to these genes (Figure 6C). The results suggest a causal relationship between the PU.1 dependent protein expression and PU.1 binding to their genes. Of note, we found that the PU.1 peak at the *IL1B* genomic region overlapped with the single nucleotide polymorphism rs1143627, which has been previously been described to be regulated by PU.1 and C/EBPβ (Zhang et al., 2014), thus confirming the notion that IL-1β regulation in TB might be via the transcription factor PU.1.

**Figure 6:**
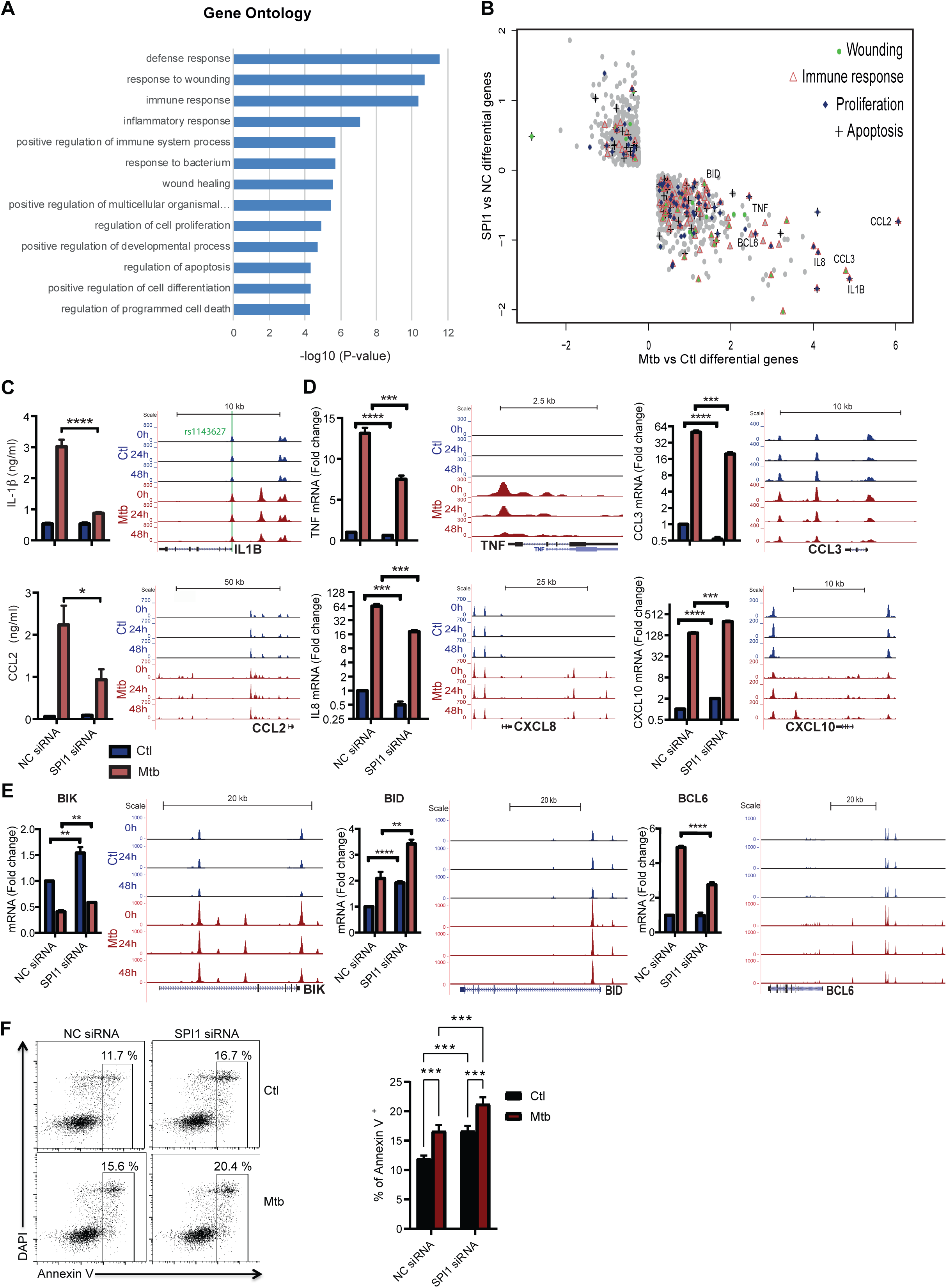
PU.1 regulates inflammatory and apoptosis response to *Mtb* infection. (A) GO analysis of PU.1 regulated genes (see also Supplemental Table 5). (B) Genes are highlighted when associated to the GO of wounding, immune response, proliferation and apoptosis. (C) Secretion of proinflammatory cytokine interleukin-1 beta (IL-1β) and chemokine monocyte chemotactic protein-1 (CCL2) by THP-1 cells in the presence or absence of PU.1 during *Mtb* infection. PU.1 binding at the promoter of each of the gene is shown on the right for control (Ctl) and infection conditions (*Mtb*). The green line in the IL-1β promoter indicates the position of the SNP rs1143627. (D) RT-qPCR experiment demonstrated the down regulation of proinflammatory cytokine (*TNF*, *IL8* and *CCL3*) and upregulation of anti-inflammatory cytokine *CXCL10* expression. The right panel indicates the corresponding PU.1 binding at each respective promoter. (E) RT-qPCR experiment revealed that the expression of proapoptotic gene *BIK* was increased in PU.1 knockdown cells; whereas antiapoptotic gene *BCL6* was decreased in PU.1 knockdown cells. Corresponding PU.1 binding to each respective promoter is shown. (F) Apoptosis induced by *Mtb* infection in THP-1 cells was enhanced in the absence of PU.1 by annexin V and DAPI staining.

Next, we validated additional PU.1 target genes by quantifying mRNA expression using qPCR (Figure 6D). *TNF*, *CCL3*, *CXCL8* and *CXCL10* were all upregulated in response to infection in a PU.1 dependent manner. Furthermore, visual inspection of PU.1 binding at the genomic locus of these genes identified multiple upregulated PU.1 peaks (Figure 6D). Since apoptosis was found in the GO results (Figure 6A), we set out to confirm the role of PU.1 in the regulation of apoptosis genes in *Mtb*-infected cells. Of note, apoptosis is an innate defense mechanism against *Mtb* infection and is inhibited by *Mtb* for its survival (Behar and Briken, 2019; Behar et al., 2011). We found the expression of proapoptotic genes *BIK* and *BID* to be upregulated whereas the *BCL6* (anti-apoptotic gene) to be downregulated in uninfected PU.1 knockdown cells (Figure 6E). *Mtb* infection alters the expression of these genes in PU.1 specific manner (Figure 6E). PU.1 binding was also altered on the genomic loci of these genes during infection. Indeed, we observed an enhanced apoptosis response in PU.1 knockdown uninfected and infected cells *viz.* increase in Annexin V^+^ cells (Figure 6F). Collectively, these data suggest that *Mtb* inhibits apoptosis response through upregulation of PU.1 to suppress proapoptotic signaling and activate prosurvival genes to maintain virulence. Taken together, these results indicate PU.1 as a critical mediator of the transcriptional response mediating inflammation and apoptosis upon *Mtb* infection.

### Pharmacological inhibition of PU.1 enhances apoptosis and restricts *Mtb* growth

Given that PU.1 knockdown enhanced apoptosis and dampened proinflammatory cytokine expression in *Mtb* infected macrophages, we next determined whether this pathway could be pharmacologically targeted. PU.1 has well-characterized DNA-binding preferences, and can be selectively inhibited using small molecules of the heterocyclic diamidine family, which disrupt PU.1 binding to DNA interfering with its transactivation activity (Antony-Debre et al., 2017). We therefore tested three of PU.1 inhibitors (DB1976 (DB19), DB2115 (DB21), and DB2313 (DB23)) for their anti-*Mtb* and pro-apoptotic activities in human primary monocyte-derived macrophages (hMDMs) (Figure 7A). All three PU.1 inhibitors were non-toxic to *Mtb*-infected hMDMs as they did not increase LDH release compared to untreated or isoniazid (INH) treated cells, which indicated the absence of membrane damage or overt cytotoxicity (Supplemental figure 2A). Only DB19 treatment (but not DB21 and DB23) led to a moderate but significant reduction in cell viability, with a ∼20% decrease in ATP-dependent luminescence relative to control (Supplemental figure 2B). We next examined the impact of PU.1 inhibition on intracellular bacterial burden using a constitutively luminescent *Mtb* strain. All three PU.1 inhibitors led to inhibition of luminescence intensity in infected hMDMs compared to untreated cells, suggesting inhibition of *Mtb* growth (Figure 7B, data from seven donors). When *Mtb* viability was directly assessed by CFU enumeration, all three PU.1 inhibitors showed significant reduction in viable *Mtb* (Figure 7C). We next determined whether PU.1 inhibition promotes apoptosis in *Mtb*-infected hMDMs. For this, we measured caspase 3/7 activity 6 days post-treatment of infected cells. All three PU.1 inhibitors increased caspase 3/7 activity relative to the infected untreated cells, with DB23 inducing the strongest and most consistent effects across donors (Figure 7D). In contrast, INH treatment did not increase caspase activity. Donor-level heatmap visualization confirmed a reproducible induction of apoptosis following DB21 and DB23 treatments (Figure 7E). DB19 and DB23 treatment of infected cells also showed a trend of reduce release of IL-1β (Figure 7E), supporting the PU.1 dependent regulation of IL-1β (Figure 6C). Together, these results indicate that PU.1 inhibition by DB21 and DB23 is well tolerated by human macrophages, significantly reducing intracellular *Mtb* burden by modulating host responses.

**Figure 7.**
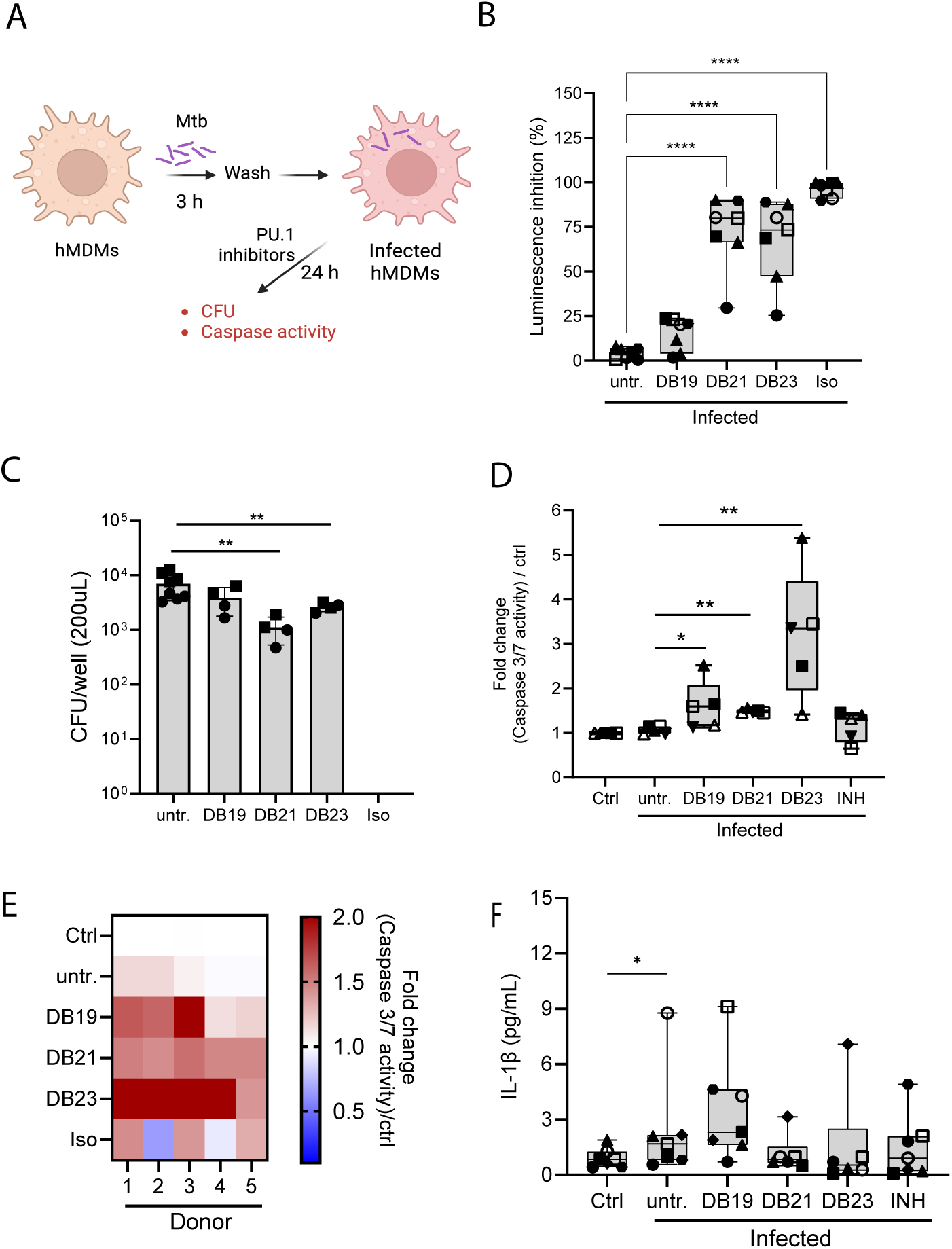
PU.1 inhibitor compounds do not induce cytotoxicity and reduce *Mtb* burden in human macrophages. (A) Schematic of infection experiment using PU.1 inhibitors. (B) Percentage inhibition of luminescence signal from *Mtb*-lux–infected hMDMs treated with PU.1 inhibitors (DB19, DB21, DB23), isoniazid (INH), or left untreated (untr.). Luminescence was measured 6 days post-infection. (C) CFU assay performed at 6 days post-infection showing bacterial load as absolute CFU per well. N=7 donors. (D) Caspase 3/7 activity measured by luminescence assay in *Mtb*-infected hMDM treated with PU.1 inhibitors, isoniazid (INH), untreated cells (untr) or uninfected cells (Ctrl). Data shown as fold change relative to uninfected control. N=5 donors. (E) Corresponding heatmap of panel D data; shows donor-wise caspase 3/7 activity fold change values relative to control. (F) Secretion of IL-1β by infected and treated hMDMs in C. Each dot in B, D and F represents an individual donor and data are shown as mean ± SD. *P*-values, one-way ANOVA with post hoc multiple comparison test; \**P* < 0.05, \*\**P* < 0.001.

## Discussion

*Mtb* modulates the host’s epigenome to promote its survival and persistence within the host (Del Rosario et al., 2022; DiNardo et al., 2020; Thomas et al., 2024). However, the critical mediator of these *Mtb*-driven epigenetic consequences remains elusive. Here, we performed genome wide H3k27ac ChIP-seq in *Mtb*-infected human cells and identified PU.1 (encoded by *SPI1* gene) as a major regulator of host-*Mtb* epigenetic network. PU.1 is considered as a pioneer epigenetic modifier in the context of hematopoiesis and immune cell development, as it can recruit chromatin remodelers and modify histone acetylation, affecting the overall structure and activity of the genome (Minderjahn et al., 2020; van Riel and Rosenbauer, 2014). Of note, *Mtb*, though primarily affects the lungs, can significantly impact hematopoiesis (Balepur and Schlossberg, 2016; Khan et al., 2020), and PU.1 has been demonstrated to directly regulate IL-1β expression that is associated with the severity of the TB disease and the treatment outcome (Zhang et al., 2014). We also show that knockdown of *SPI1* gene dampens *Mtb*-driven IL-1β expression (Figure 6C). Data from *in vitro* and TB patient monocytes indicate that PU.1 gene and protein expression might be up-regulated during *Mtb* infection likely via the activation of the *SPI1* super enhancer. This, up-regulation of PU.1 was beneficial to the survival of the intracellular bacteria.

Direct evidence of the role of PU.1 in infectious disease is limited. It has been shown that Epstein-Barr virus infection modulates transcription program in B cells through PU.1, suppressing host defense pathways to achieve immortal cell growth (Lin et al., 2015; Zhao et al., 2011). Mouse alveolar macrophages infected with *Pneumocystis* have decreased expression of PU.1, which leads to the downregulation of macrophage phagocytic receptor Dectin-1 (Zhang et al., 2010). Furthermore, PU.1 can shape mouse neutrophil defence program and exorbitant inflammatory responses by broadly inhibiting the accessibility of enhancers via the recruitment of HDAC1 (Fischer et al., 2019). These studies along with ours thus provide pieces of information that PU.1 indeed plays a key role in infectious disease. An important question to be addressed remains, the mechanisms that control the level of PU.1 in the infected host.

Our genetic and pharmacological investigation demonstrate modulation of apoptosis and inflammation as the downstream effect of inhibiting PU.1 (Figure 6 and 7). PU.1 can both promote and inhibit apoptosis depending on the cellular context. It can induce apoptosis by upregulating pro-apoptotic genes like TRAIL and DR5, or it can promote cell survival by inhibiting apoptotic pathways, such as NF-κB (Haimovici et al., 2017; Yuki et al., 2013). In preleukemic cells, PU.1 promotes resistance to apoptosis by epigenetic silencing of the proapoptotic factor BIM through recruiting PRC2 repressive complex and promoting the trimethylation of H3K27 on the promoter (Ridinger-Saison et al., 2013). We found enrichment of PU.1 binding on key gene locus regulating apoptosis, thus suppressing PU.1 might affect the epigenetic programming at these loci resulting in modulation of respective gene expression (Figure 6E). Hence, it is plausible that *Mtb* inhibits apoptosis by epigenetic control of gene expression through manipulating the interaction between PU.1 and other cofactors, thus disrupting the balance of pro- and anti-apoptosis molecules. PU.1 inhibitor experiments in hMDMs further support this balance (Figure 7).

This study was conducted entirely on *in vitro* human infected cells, and therefore the relevance of PU.1 inhibition during *in vivo* infection remains to be established. Critical aspects such as compound bioavailability, systemic toxicity, and efficacy in reducing bacterial burden and pathology in lung tissue require validation in preclinical mouse models. However, mouse studies may not always accurately reflect human cell behavior *in vitro* (Masopust et al., 2017). Although we have employed three structurally related PU.1 inhibitors with known specificity, potential off-target effects or compound-specific activities cannot be fully ruled out. We also did not assess the epigenetic network following PU.1 inhibition by small molecules, which could provide critical insights into the immunomodulatory consequences of targeting this transcription factor. Despite these limitations, our findings identify PU.1 as a critical epigenetic regulator that modulates the balance between macrophage response and bacterial survival during *Mtb* infection. Targeting PU.1 enhances macrophage-intrinsic antibacterial responses without broad cytotoxicity, which support the concept of PU.1 inhibition to induce macrophage anti-bacterial pathways leading to the design of a host-directed therapeutic strategy for TB (Sweet et al., 2025; Wallis and Hafner, 2015).

## Material and Methods

### Cell culture

Human monocytic leukemia cell line THP-1 was maintained in RPMI-1640 medium (Gibco) supplemented with 10 % fetal bovine serum, 2 mM glutamine, 1 mM sodium pyruvate, 1 % non-essential amino acids, and 100 μg/ml kanamycin, hereafter referred as complete medium. The culture flask was placed in a moist incubator at 37 °C supplied with 5% CO_2_.

### Preparation of human monocyte derived macrophages

To prepare human monocyte-derived macrophages (hMDM), CD14^+^ monocytes were isolated directly from buffy coat (Etablissement Français du Sang, convention N° 21RB2025-028-R) using magnetic positive selection (Miltenyi Biotec, #130-114-976), following the manufacturer’s instructions. The purified monocytes were differentiated into macrophages by culturing them in complete RPMI-1640 medium supplemented with 10% FCS (Sigma-Aldrich, #F7524), and 100 ng/ml recombinant human M-CSF (R&D systems, #216-MC-500) for 6 days. The differentiated macrophages were collected and cultured in fresh medium without M-CSF overnight prior to infection.

### RNA interference

Knockdown of PU.1 in THP-1 cells was achieved by transfection of pooled DsiRNA (Integrated DNA technologies, #HSC.RNAI.N003120.12.1, #HSC.RNAI.N003120.12.2 and #HSC.RNAI.N003120.12.3) using Lipofectamine RNAiMAX (Invitrogen, #13778-150) according to the manufacturer’s protocol. For one well of the 6-well plate, 5 x 10^5^ THP-1 cells were seeded one day before transfection in antibiotic-free RPMI-1640. Three μl of DsiRNA duplex (45 pmol) was diluted in 200 μl of RPMI-1640, meanwhile 9 μl of RNAiMAX was diluted in 200 μl of RPMI-1640. The diluted DsiRNA was mixed with diluted RNAiMAX and incubated at room temperature for 15 min, then the complex was added to cells drop-wisely. Knockdown of PU.1 in hMDM cells was achieved by electroporation of pooled DsiRNA using Amaxa^®^ nucleofector machine with Cell Line Nucleofector^®^ Kit V (Lonza, #VCA-1003) and program U-001. Thirty-six hours post-transfection, cells were infected with *Mtb* or BCG, and followed by downstream analysis.

### Bacterial strains and growth conditions

*Mtb* H37Rv and *Mycobacterium bovis* BCG strain was cultured as described previously (Cheng et al., 2017). Briefly, *Mtb* H37Rv and BCG were cultured in Middlebrook 7H9 broth (BBL Microbiology Systems, USA) supplemented with 10% Middlebrook oleic acid, albumin, dextrose, and catalase enrichment (Difco laboratories, USA) and 0.05% Tween 80 at 37°C to an optical density of 0.4-0.5 (OD_600_). The strain H37Rv::lux was constructed by Dr. Claud Gutierrez (CNRS – IPBS) after transformation of *Mtb* wild type strain H37Rv with the integrative plasmid pMV30-hsp-Lux13 (Addgene #26161) and cultivated with the same medium described before plus kanamycin (50µg/mL). Logarithmically growing mycobacteria were harvested, resuspended in fresh 7H9 broth with 20% glycerol and frozen in −80°C in small aliquots. Representative vials were thawed and enumerated for viable CFU on Middlebrook 7H11 plates.

### *In vitro* mycobacterial infection and enumeration

THP-1 and hMDM were infected with *Mtb* or BCG at multiplicity of infection of 5. Briefly, frozen mycobacteria were thawed, washed and resuspended in plain RPMI-1640 medium at concentration of 3×10^7^ / ml. THP-1 and hMDM were seeded in 6-well plate at concentration of 3×10^6^ / ml in complete RPMI-1640 without antibiotics. Concentrated mycobacteria were added into THP-1 and hMDM culture and incubated at 37 °C for 3 h. Infected cells were collected and washed twice to remove extracellular mycobacteria, and seeded in triplicates for downstream experiments. This time point was referred as time 0 h post-infection. To assess intracellular mycobacterial proliferation, at indicated time points post-infection, infected THP-1 and hMDM were harvested, washed twice with PBS, and then lysed by resuspension in 0.6 % SDS solution with vigorous pipetting. Cell lysates were 10x serial diluted in 7H9 broth and plated on Middlebrook 7H11 agar on quadrant plates in triplicates. The plates were incubated at 37°C for 3 weeks before colonies were counted visually. Colony forming unit (CFU) obtained from two dilutions were used to calculate the total number of CFU/ml.

### ChIP-seq of histone H3K27ac and PU.1

ChIP-seq was performed as we described before (del Rosario et al., 2015). Briefly, 5 million cells were cross-linked with 1% formaldehyde for 15 minutes and excess formaldehyde quenched by addition of glycine (0.625 M). Nuclei were prepared and collected under standard procedures and re-suspended in 300 μl SDS lysis buffer (1% SDS, 1% Triton X 100, 2 mM EDTA, 50 mM Hepes-KOH [pH 7.5], 0.1% Na dodecyl-sulfate, Roche 1X Complete protease inhibitor). Nuclei were lysed for 15 minutes, after which sonication was used to fragment chromatin to an average size of 200–500 bp (Bioruptor Next gen, Diagenode). Nuclear lysates were prepared following standard procedures. Protein-DNA complexes were immuno-precipitated using 3 μg of H3K27acetyl antibody (Actif motif, #39133) or 10ug of PU.1 antibody coupled to 50μl protein G Dynal beads (Invitrogen) overnight. The beads were washed, and protein-DNA complexes were eluted with 150 μl of elution buffer (1% SDS, 10 mM EDTA, 50 mM Tris.HCl [pH 8]), followed by protease treatment and de-crosslinking at 65°C overnight. After phenol/chloroform extraction, DNA was purified by ethanol precipitation. Library preparation was performed followed by multiplexed 36bp sequencing on an Illumina HIseq (del Rosario et al., 2015).

### Read alignment and peak calling

Reads were mapped to the human genome (hg19) using BWA. Duplicate reads were filtered out using SAMtools and only reads with mapping quality score ≥ 10 were used for downstream analysis. Peaks were called using DFilter (Sun et al., 2016). On average 17,770 peaks were called with the *P*-value threshold of 1e-6 for H3K27ac libraries. Then top 5K peaks of each H3K27ac library were selected to form a union of peak regions (7423 regions) for all libraries. The union of peak regions is defined as any peak regions from at least two libraries. For each PU.1 ChIP-seq library, approximately 40,000 peaks were called with the *P*-value threshold of 1e-10. 52314 union PU.1 peak regions were selected subsequently.

### Peak height normalization

Read counts in 100 bp bin of each library were first normalized against the input. Then read counts were adjusted by normalizing their GC-content against the average GC-content of all libraries. In each union peak region, the total number of read counts after GC normalization was defined as the peak height. To remove the variability across libraries, quantile normalization was performed subsequently for the peak heights in the union peak region.

### Clustering

The log2 fold change of each time point was taken (*Mtb*/ control) on the union of the top 5K peaks from above. K-means clustering was then performed using Cluster 3.0 with K = 7 and visualized the resulting matrix using Java TreeView. To remove noise, we defined variation of peak height that is below log2 of 1 and log2 of −1 as 0. The peak locations from each cluster were then analyzed for Gene ontology enrichment using GREAT (McLean et al., 2010) using the default settings, with the whole data set as background.

### Motif analysis

Motif analysis was performed on the peak regions of each cluster independently, using findMotifsGenome.pl code from the Homer ChIP-seq pipeline (Heinz et al., 2010). This was performed using the mknown setting with the Transfac vertebrate database as the set of known motifs. In addition, candidate TF motifs were kept if the fold change was above 2 and a *Q*-value ≤ 0.01 after multiple hypothesis correction using Benjamini Hochberg method. Motifs were associated with the genes using GSEA and their respective family and subfamily according to GeneXplain TF classification.

### Microarray analysis

Microarray (Illumina, HumanHT-12v4) was performed in triplicates and genes with an average signal of < 100 were removed resulting in a subset of 9557 genes. We then defined *Mtb* differentially expressed genes as genes that had log2 (fold-change) greater than 0.2 for the *Mtb* infected vs control samples and that were significantly differentially expressed. The significance of differential expression was assessed using a Student *t*-test followed by Benjamini-Hochberg FDR correction for multiple testing: FDR< 10%. Next, we calculated the expression fold-change upon PU.1 knockdown at 24h of *Mtb* infection (targeting vs control siRNA). We then defined PU.1 regulated genes (940) as those that were either upregulated upon *Mtb* infection (see above) and downregulated (Student *t*-test < 0.05) by PU.1 knockdown or downregulated by *Mtb* and upregulated (Student *t*-test < 0.05) by PU.1 knockdown. Gene Ontology enrichment analysis was performed on PU.1-regulated genes with all expressed genes as the background set using DAVID (Huang da et al., 2009). Genes directly regulated by PU.1 were defined as those that were PU.1-regulated by the above criterion and had a PU.1 ChIP-seq peak within 5,000 basepairs of the transcription start site.

### Super enhancer analysis

We use the strategy defined in to identify super enhancers (Pott and Lieb, 2015). Briefly, constitutive enhancers regions within 12.5kb of each other are combined into a single domain, these domains were sorted in ascending order and then plotted according to the ChIP-seq peak height. The single domain that were above the point of inflection of the curve were defined as super-enhancers (Del Rosario et al., 2022; Hnisz et al., 2013).

### Real-time reverse transcription PCR

Total RNA was extracted using RNeasy Mini Kit (QIAGEN, #74106) from control and mycobacterial infected cells at indicated time points. Five micrograms of total RNA were converted to cDNA using the iScript^TM^ Advanced cDNA Synthesis Kit (Bio-Rad, #172-5037). Real-time PCR was performed using SsoAdvanced^TM^ SYBR^®^ Green Supermix (Bio-Rad, #172-5261). The relative quantities of the gene tested per sample were calculated against RPL13A using the delta delta threshold cycle (ΔΔCt) formula.

### Cell lysate preparation and western blot

THP-1 and hMDM cells were harvested and lysed in RIPA buffer (Sigma-Aldrich, #R0278), supplemented with protease inhibitors (Roche, #05892970001) and phosphatase inhibitors (Roche, #04906837001) at 2 x 10^7^ cells/ml. The lysate was incubated on ice for 10 min and cell debris was removed by centrifugation at 14000 rpm for 10 min at 4°C. The protein concentration was determined using the BCA protein assay kit (Pierce, #23225) by measuring absorbance at 562 nm compared to a protein standard. Cell lysate (20 ug of protein) was resolved on SDS-PAGE and transferred to PVDF membranes. The following antibodies were used: PU.1 and GAPDH (Cell Signaling Technology, #2258 and #2118).

### Apoptosis assay

Early apoptosis in THP-1 cells was detected by Annexin V staining. Briefly, THP-1 cells were transfected with non-targeting control (Integrated DNA technologies, DS ScrambledNeg) or PU.1 DsiRNAs for 36 h, and infected with *Mtb*. Twenty-four hours post-infection, cells were stained with Annexin V-FITC and propidium iodide following the manufacturer’s protocol (Cell Signaling Technology, #6592). Samples were acquired using the Miltenyi cytometer installed with the MACSQuantify software, and data was analyzed using FlowJo software.

Caspase activities in BCG-infected THP-1 cells were measured using the Caspase-Glo^®^ 3/7 assay (Promega, #G8090). THP-1 cells were transfected with non-targeting control or PU.1 DsiRNAs for 36 h and infected with BCG. Twenty-four hours post-infection, cells were seeded into 96-well plate at 20000 cells per well in triplicates. Caspase-Glo^®^ 3/7 reagent was added to wells, and luminescence was recorded at 5 min intervals during a one-hour period. Background readings were determined from wells containing culture medium without cells and were subtracted from signal readings. Results were plotted as luminescence signals in each sample over a one-hour period.

Caspase activity in *Mtb*-infected hMDM cells was quantified by measuring caspase 3/7 activity using a luminescence-based assay. Luminescence was measured in whole wells 6 days after infection and treatment. Results were normalized to uninfected controls and reported as fold change.

### Cytokine measurement by ELISA

Detection of human IL-1β and MCP-1 in the supernatant of THP-1 cells was performed using the following antibody pairs: IL-1β (Biolegend, 508202, 508302), MCP-1 (Biolegend, 505906, 502609). Each sample was measured in triplicates, and the average value with standard deviation was plotted.

### Human lymph node TB immunostaining

Paraffin-embedded cervical lymph node tissue sections (5 µm) from extrapulmonary TB patients were deparaffinized in xylene and rehydrated through a graded ethanol series. Endogenous peroxidase activity was quenched with 0.3% hydrogen peroxide in PBS for 10 minutes. Following antigen retrieval in citrate buffer (pH 6.0) at 95°C for 20 minutes, sections were blocked with 5% normal goat serum and incubated overnight at 4°C with a rabbit monoclonal anti-PU.1 antibody (Cell Signaling Technology, clone 2258, 1:200). After washing, slides were incubated with an HRP-conjugated secondary antibody and developed using DAB (DAKO) according to the manufacturer’s instructions. Hematoxylin was used for counterstaining. Slides were examined by light microscopy to evaluate nuclear PU.1 expression in macrophages, including epithelioid cells and multinucleated giant cells.

### Macaque lung immunostaining

Lung tissues from naïve or BCG vaccinated cynomlgus macaques (Macaca fascicularis) that were subsequently aerosol infected with 1000 CFU of *Mtb* strain CDC1551 (Kaushal *et al*., 2015) were formalin-fixed, paraffin-embedded, and sectioned at 5 µm thickness. After deparaffinization and rehydration, antigen retrieval was performed using citrate buffer (pH 6.0) at 95°C for 20 minutes. Sections were blocked in 5% normal donkey serum and incubated overnight at 4°C with primary antibodies: rabbit anti-PU.1 (CST, 1:200) and mouse anti-CD163 (Abcam, #ab182422, 1:100) or anti-CD68 (Bio-Rad, #MCA1957, 1:100). Fluorescent secondary antibodies (donkey anti-rabbit Alexa Fluor 488 and donkey anti-mouse Alexa Fluor 594; Invitrogen) were applied for 1 hour at room temperature. Nuclei were stained with DAPI. Sections were mounted with antifade reagent and imaged using a Leica TCS SP8 confocal microscope. Quantification of PU.1^+^CD68^+^CD163^+^ double-positive cells was performed in multiple fields per animal using ImageJ.

### PU.1 inhibitor compounds

Three heterocyclic diamidine compounds—DB1976, DB2115, and DB2313—were used to pharmacologically inhibit PU.1 activity. These small molecules were originally developed and characterized by Antony-Debré et al. (Antony-Debré *et al*., 2017) as allosteric inhibitors of PU.1 that disrupt DNA binding by targeting the AT-rich minor groove adjacent to PU.1 consensus motifs. Compounds were obtained from Sigma-Aldrich and reconstituted in sterile DMSO. Working concentrations were selected based on prior IC data and validated in preliminary toxicity and efficacy assays.

### PU.1 inhibitor treatment of hMDM

Human MDMs were infected with *Mtb* H37Rvlux strain at a MOI of 1 for 4 hours, followed by washing to remove extracellular bacteria. Infected and uninfected hMDMs were then treated with 1 µM of PU.1 inhibitors (DB19, DB21, DB23), or a positive control isoniazid (INH) or left untreated in fresh medium. Inhibitor concentrations were kept constant across experiments. Assays were performed at 6 days post-treatment.

### LDH release assay

LDH release was measured to evaluate potential PU.1 inhibitor cytotoxicity on hMDMs. Cell culture supernatants were collected 6 days after treatment. LDH activity was assessed using the CyQUANT™ LDH Cytotoxicity Assay Kit (Invitrogen, # 16280972). Total lysis controls were generated by incubating cells with lysis buffer (0.1% Triton/water). LDH activity in experimental wells was normalized to total lysis and expressed as percentage of maximum release.

### Cell viability assay

Cell viability was determined by measuring total ATP content in cell lysates using a luminescence-based readout (Promega, # G9242). Signal intensity was used as a proxy for viable cell number. Values were normalized to uninfected control cells.

### Luminescence assay for *Mtb* load in hMDMs

To assess bacterial burden, luminescence was measured directly from infected macrophages using a constitutively luminescent *Mtb* strain. Luminescence values were recorded 6 days post-treatment and expressed as percentage inhibition relative to untreated infected controls.

### CFU enumeration assay

To determine viable intracellular *Mtb*, H37Rvlux infected cells were lysed 6 days post-treatment and serial dilutions of the lysates were plated on 7H11 agar. Plates were incubated at 37°C for 3–4 weeks before counting colonies. Results were expressed as colony-forming units (CFU) per well and normalized to the untreated condition.

## Acknowledgements

We acknowledge the GIS sequencing platform for sequencing the ChIP-seq libraries. We acknowledge the SIGN transcriptomics platform for carrying out the microarray experiments. We also thank Claude Gutierrez for kindly providing the *Mtb* H37Rv::lux strain. J.P and C.C are supported by the ANRS-MIE Project Riposte. CC is supported by the European Commission, Project ITHEMYC “Novel Immunotherapies for tuberculosis and other mycobacterial diseases”, Grant agreement n° 101080462. A.S is supported by A*STAR ID Labs, BMRC CDA award, MOH-OFIRG23jul-0016 and NIH 1R01HL152078.

## Author contributions

J.P, L.J, S.P, G.D.L and A.S conceived and designed the study. L.J performed all *in vitro* experiments. T.W.F and D.K generated and analyzed monkey data. CO.C and G.P collected and analyzed human image data. C.B and CE.C performed PU.1 inhibitor experiments. J.P, W.S & S.P generated epigenetic data. L.J and A.S generated transcriptomics data. J.P, L.J, W.S, A.M and A.S performed the analysis. J.P, L.J & AS wrote the paper with help from all co-authors.

