## Supplementary figures and images for "PU.1 Shapes Host Epigenetic Responses to Mycobacterium tuberculosis and Represents a Target for Host Directed Therapy"

### Supplemental Figure 1

**Supplemental figure 1**

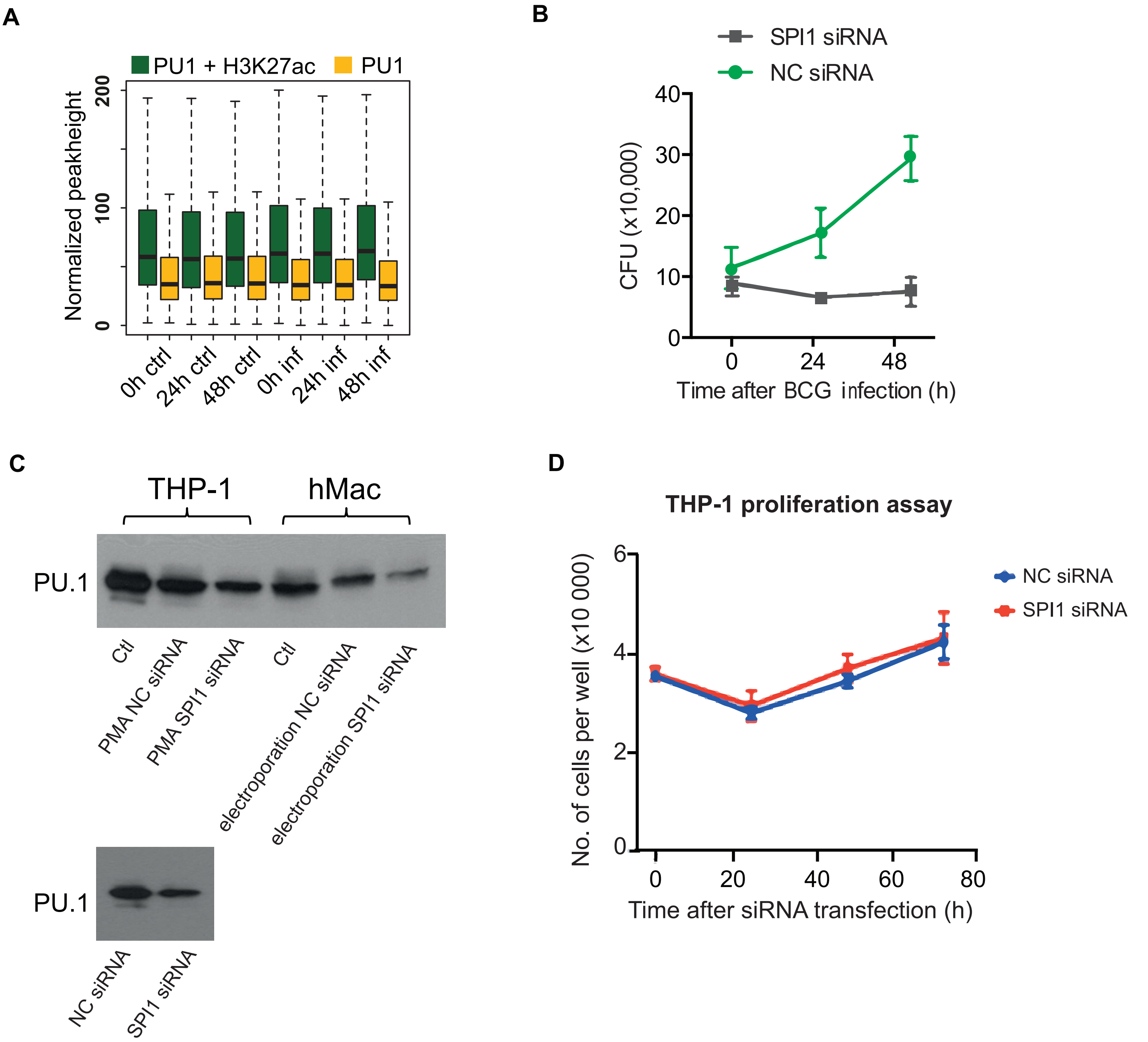

### Supplemental Figure 2

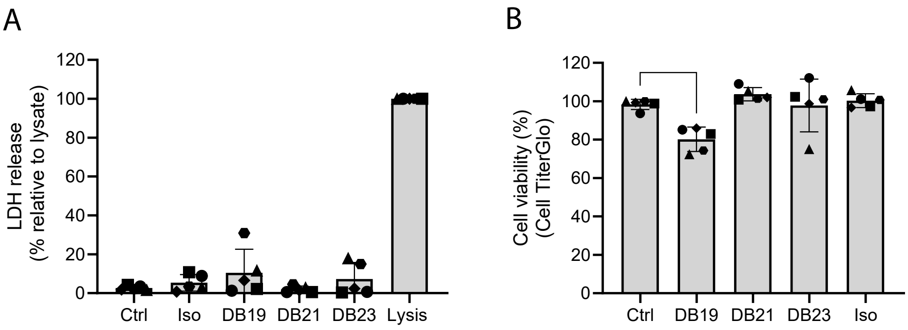
