## Supplemental table 4 for "PU.1 Shapes Host Epigenetic Responses to Mycobacterium tuberculosis and Represents a Target for Host Directed Therapy"

**Table S1**. TB datasets used in this study.

| **ID** | **Country of cohort** | **Healthy controls** | **Active TB** | **Post-treatment active TB** | **Latent TB** | **Other diseases** | **Study reference No.** | **Microarray platform** | **GEO accession** |
| --- | --- | --- | --- | --- | --- | --- | --- | --- | --- |
| UK ‘10 | United kingdom | 12 | 21 | - | 21 | - | 26 | Illumina Human HT-12 v3.0 | GSE19444 |
| SA’ 10 | South Africa | - | 20 | - | 31 | - | 26 | Illumina Human HT-12 v3.0 | GSE19442 |
| SA ‘12 | South Africa | - | 29 | 29 | 38 | - | 35 | Illumina Human HT-12 v4.0 | GSE40553 |
| SA ‘13 | Malawi | - | 97 | - | 97 | 83 | 40 | Illumina Human HT-12 V4.0 | GSE37250 |
