## Supplemental Figure Legend for "PU.1 Shapes Host Epigenetic Responses to Mycobacterium tuberculosis and Represents a Target for Host Directed Therapy"

**Supplemental Figures**

**Supplemental Figure 1.**
**A.** ChIP-seq normalized peak height of H3K27ac + PU.1 peaks (overlapped peaks) and PU.1 peaks which do not overlap with H3K27ac peaks at multiple time points (0h, 24h, 48h) after MtB infection in THP-1 cells.
**B.** Bacterial burden measured as colony-forming units (CFU) over time (0h, 24h, 48h) after BCG infection in SPI1 knockdown cells vs siRNA control.
**C.** PU.1 protein levels in THP-1 and human macrophages (hMac) treated with NC siRNA or SPI1 siRNA, with and without PMA stimulation, assessed by Western blot.
**D.** THP-1 proliferation assay over time following transfection with NC or SPI1 siRNA, quantified by cell count.

**Supplemental Figure 2.**
**A.** Cytotoxicity measured by LDH release in THP-1 cells treated with control (Ctrl), isotype control (Iso), and PU.1 inhibitors DB19, DB21, and DB23. Values are expressed as % relative to total lysis.
**B.** Cell viability of THP-1 cells treated as in A, assessed by CellTiter-Glo assay. Values are expressed as % viability relative to control.

**Supplemental Tables**

**Supplemental Table 1.** PU.1 and H3K27ac ChIP-seq datasets.

- *Sheet “Ac_PU1_genes_clusters”*: Annotated H3K27ac peaks and their nearest genes (n=7,846), clustered based on dynamic changes at 0h, 24h, 48h, and 72h post-infection, with corresponding PU.1 binding status.
- *Sheet “all_PU1_peaks”*: Full list of PU.1 peak regions (n=66,925) with read counts at multiple time points.
- *Sheet “all_AC_peaks”*: All detected H3K27ac peaks (n=45,441) with signal intensities across time points in control and infected conditions.

**Supplemental Table 2.** Gene Ontology enrichment analysis for H3K27ac-PU.1 clusters.

- *Sheet “all”*: Summary of 16 enriched ontology terms across all clusters.
- *Sheets “cluster3” to “cluster7”*: Cluster-specific enrichment for biological processes, including adjusted p-values, enrichment scores, and foreground gene lists.

**Supplemental Table 3.** De novo motif analysis of cluster-specific PU.1-H3K27ac peaks.

- *Sheets “Cluster 3” to “Cluster 7”*: Enriched transcription factor motifs with consensus sequences, p-values, fold enrichment, and annotation of gene family and subfamily.

**Supplemental Table 4.** Public tuberculosis (TB) transcriptomic datasets used in the study.

- Lists cohort metadata from four studies (UK 2010, SA 2010, SA 2012, Malawi 2013), including disease groups, microarray platforms, and GEO accession numbers.

**Supplemental Table 5.** Gene expression and differential analysis from siRNA knockdown and MTB infection experiments.

- *Sheet “Expression file”*: Raw expression matrix for 47,323 probes across 15 experimental conditions (KD PU.1 +/- and MTB +/- infection).
- *Sheet “SE genes”*: List of 453 super-enhancer-associated genes with genomic coordinates and distance to TSS.
- *Sheet “PU1 binding red”*: PU.1 binding sites reduced upon SPI1 knockdown (n=9,730), annotated with gene proximity.
- *Sheet “GO”*: Gene Ontology enrichment terms derived from PU.1-regulated gene sets.
- *Sheet “Patient DE genes green”*: 1,979 genes differentially expressed in patients vs controls with fold change and statistical metrics.
- *Sheet “MTB Inf DEG”*: Differentially expressed genes (n=5,741) in response to MTB infection.
- *Sheet “PU1KD DEG”*: Differential gene expression (n=4,105) following SPI1 knockdown, including log fold change and adjusted p-values.
